# HI-JEPA: A World Model of Molecular Organization Learned from Measured Proximity

**DOI:** 10.64898/2026.08.14.744901

**Authors:** Ryan Shihabi, Siddhant Karmali, Brent Vaughan, Sharief Taraman, Manolis Kellis

## Abstract

Proteins act through the company they keep. Which molecules occupy the same nanoscale neighborhood in intact tissue determines what can physically interact, and disease rearranges those neighborhoods before it changes anything a sequence records. That quantity (measured proximity between molecular species in unperturbed tissue) has never been acquired broadly enough to train on. Published colocalization arrives study by study and never accumulates into a graph. The measurement has to be made rather than collected.

We built ASCEND, a spatial computing platform that measures pairwise molecular proximity from expansion microscopy at molecular resolution in intact tissue, and applied it to 164 proteins across 37 imaged regions in five studies, spanning cultured neurons, isolated synapses and mouse cortex in disease and control. HI-JEPA is a representation trained on those measurements. Each protein is one embedding, trained to predict the embeddings of its measured neighbors in latent space; it never reconstructs its input and generates no negatives. A set of proteins measured in one neighborhood forms a *configuration*, which is the object the model perturbs and plans over.

The representation performs operations a sequence model cannot. It names a protein from the bare geometry of a microscopy point cloud, matched against 234,048 deposited structures, at top-1 accuracy 0.748 against a chance rate of 1.0 × 10^-5^. It predicts physical interaction between sequence-dissimilar proteins that were both withheld from training at AUC 0.908, where ESM-C 6B reaches 0.514 against partner-count-matched negatives. It recovers a held-out complex member in the top 100 of 13,447 candidates at recall 0.954, against 0.514 for a ranking built from complex frequency alone. Asked which partners a knockout disrupts, it recovers the experimentally observed ones at recall@100 0.640; asked the same question about a different protein, with the ranking rule and denominators unchanged, it recovers 0.028, so the answer follows the action.

Given 5xFAD mouse cortex with no disease label, no reward and no indication that amyloid is relevant, ranking 1,574 measured assemblies by their departure from wild type returns amyloid-*β* bound to AMPA receptor subunits in nine of the top ten. Planning over the same configurations independently selects the same subunits (GluA2, GluA3, GluA4) and predicts that disrupting the PSD-95 scaffold worsens the configuration, both agreeing in sign with experiments the model never saw. Ablating the measured-proximity channel at training time degrades cross-scale partner recovery from median rank 14 to 68 while leaving navigation and within-scale dynamics intact; ablating the perturbation channel does the reverse. The cross-scale capability therefore comes from the measurement and not from having seen more data.

The intended application is target nomination in diseases where sequence and structure supply no starting point.

**Note:** This is a capability report. The architecture, the training procedure and the acquisition protocol are proprietary and are not described. Section 4.2 gives the evaluation protocol behind every number reported.

## 1. Introduction

### 1.1 The Quantity That Is Missing

Whether two proteins can interact is constrained before any chemistry: they have to be in the same place. In intact tissue, which molecules occupy a shared nanoscale neighborhood is a physical fact about the sample, and it is not recoverable from the molecules themselves. Sequence records evolutionary history. A solved structure records where atoms sit inside one molecule, and a solved complex records a pair already known to associate, a different quantity from which molecules sit together in a tissue that nobody has perturbed. Expression records how much of something is present, not where.

Disease acts on this layer directly. Amyloid-*β* does not change the sequence of an AMPA receptor; it changes what that receptor sits next to. A pocket that opens only in a pathological conformation, or an interface that forms only because a disease state has brought two proteins into contact, is invisible to any method reading one molecule at a time.

The obstacle is not imaging. Public repositories already hold more microscopy than can be processed and the volume continues to grow. What is missing is the step that turns a scan into a measurement: which molecules occupied one neighborhood, at what separation, across enough proteins to form a graph rather than a finding. Published colocalization arrives study by study and never accumulates into that graph. The measurement has to be made rather than collected.

**A necessary caveat:** Measured proximity is not binding. Two proteins can share a neighborhood without contact, and at the resolutions reported here proximity is a necessary but not sufficient condition for physical interaction. Throughout this paper, results are labeled by what grounds them: *proximity-grounded* where the readout is colocalization or co-assembly (Sections 6.2, 8.2, 8.4, 11.2), and *interaction-grounded* where the readout is scored against experimentally documented physical interactions or curated complexes (Sections 6.1, 7.1, 7.2). The claim of the paper is that a representation trained on the first transfers to the second.

### 1.2 Approach

We train on measured molecular proximity acquired at nanoscale resolution, where one acquisition spans co-assembly through tissue-scale organization and resolves tens of protein species at once. Measured distances become proximity relations (Section 4.1). A representation is then trained so that each protein’s embedding predicts the embeddings of its measured neighbors in latent space, without reconstructing the input and without generated negatives (Section 5).

Because every protein occupies one embedding space, a set of proteins measured together is a configuration in that space, and a relation crossing between scales is a distance in one coordinate system rather than a mapping between separate models. That is what the paper means by *hierarchical*.

### 1.3 Relation to Other Uses of the Term “World Model”

Three methodologies presently claim the term in biology. The first scales a language model over amino-acid sequence until its representation organizes in agreement with established biology, and supplies the general-purpose embeddings most of computational biology runs on [10]. The second simulates a cell, learning how an expression profile responds to a perturbation [12, 13, 14]. The third wraps a language model around analysis tooling and treats the chain of invocations as a model of the system.

A world model, as originally defined, has three operations: it turns an observation into a state, predicts how that state changes under an action, and scores the action by whether the state moved toward a goal [4, 5, 15]. Each line of work satisfies a different subset. A sequence embedding holds no configuration to perturb, so its transition is undefined. A tool-calling system answers by chaining invocations, so the answer depends on which tools ran and in what order, and what it holds afterwards is text about biology. Virtual-cell programs do define a state and a transition, scoped to a single cell, with the state grounded in the geometric similarity of pretrained embeddings over curated relationships. HI-JEPA’s state is a measurement.

### 1.4 Contributions

· **Measured proximity as the learning signal.** The supervisory relation is a physical measurement rather than an inference from sequence similarity or a curated annotation (Sections 3, 4).
· **Prediction from positives alone.** A non-contrastive predictive objective exceeds a contrastive control trained on identical data across navigation, interaction, function and cross-scale retrieval, with no dimensional collapse and no generated negatives, at roughly 2,000× fewer pair comparisons (Section 5).
· **One state across scales.** A single latent carries every result from compound binding through complex and cellular to tissue (Section 10).
· **Grounding tested by removal.** Removing the measured-proximity relation degrades cross-scale partner recovery while leaving representation quality and within-scale dynamics intact; removing the perturbation relation does the reverse (Section 9).
· **A worked disease case.** From tissue alone, with no disease label, the model recovers the assemblies independent experiments identify as the mediators of amyloid pathology (Section 8).

## 2. Related Work

HI-JEPA belongs to the family of joint-embedding predictive architectures, which learn by predicting the latent representation of one view from another without reconstructing the input. This lineage includes image-based predictive models [9] and the broader class of non-contrastive self-supervised methods built on redundancy-reduction [7] and self-distillation [8] objectives, which avoid explicit negative sampling. A recent line provides distributional guarantees on the embedding marginal through a sketched isotropic-Gaussian regularizer [6]. We adopt this family for a biological relation graph in which the predicted neighbors are distinct related entities rather than augmentations of a single input, and we select the collapse-prevention objective empirically for that setting (Section 5).

The world-model formulation originates outside biology [4, 5], where a state evolves under actions as *z*^′^ *= F(z, a)* and supports simulation and planning over trajectories. We adopt those criteria directly, following recent critiques of how the term is used [15].

Sequence- and structure-based foundation models [10, 11] provide the prevailing infrastructure for protein representation and serve as the comparison baselines of Section 6.1. A folding model can be read as a transition, where the action is an edit to the sequence and the state is the fold of one molecule. The perturbations acted on here admit no such edit: removing a constituent from an assembly, or moving a tissue between disease states, changes a configuration of many molecules rather than the shape of one.

Virtual-cell programs [12, 13, 14] take the cell as the atomic unit, where HI-JEPA models the measured physical organization of molecules across scales. Predictive perturbation models [16] capture one projection of cellular response within a fixed observation space and are complementary to this work.

## 3. The Model

### 3.1 Terms

· **Embedding**, the single latent vector assigned to one protein.
· **Configuration**, the set of embeddings of a set of proteins measured together; equivalently, the world-model **state**. The two words are used interchangeably below only where “state” carries the world-model sense explicitly.
· **Action**, a perturbation applied to a configuration: removal or stabilization of a constituent, or a modeled intervention.
· **Relation**, a measured or curated pairing between two proteins that supplies training signal. Each relation type is a **channel**; channels are what Section 9 ablates.

### 3.2 Construction

We built one model spanning local structural morphology, relational interactions between molecular neighbors, and regional tissue state. Interaction-scale relations are learned from measured spatial proximity, and the regional state summarizes the molecular interactions within a region. This layering places the molecular and tissue scales in one coordinate system, so a relation crossing between them is a distance in that space rather than a mapping between separate models (Section 9).

Each protein is one embedding, informed by whichever measured and reference channels exist for it (Section 4.1). The embedding is trained so that a protein’s vector predicts the vectors of its measured neighbors. One representation then supports retrieval, completion of a partial configuration, scoring of candidate relations, and identification of a measured configuration against the state.

An action is applied to the configuration and the model predicts what follows. A learned operator does this in both measurement spaces, and it is trained on measured perturbation response (a set-structured transformer over the control-cell population, conditioned on the action as an input token and predicting a residual change; the action is the model’s own latent for the perturbed gene; trained on published single-gene perturbation response across four cell lines and evaluated on 2,749 held-out gene-and-cell-line combinations). Reward is displacement along a measured axis between a diseased and a healthy configuration, and planning searches action sequences that maximize it (greedy over single interventions, horizon 8 with early termination, action space 28) (Sections 7, 8).

**Figure 1.**
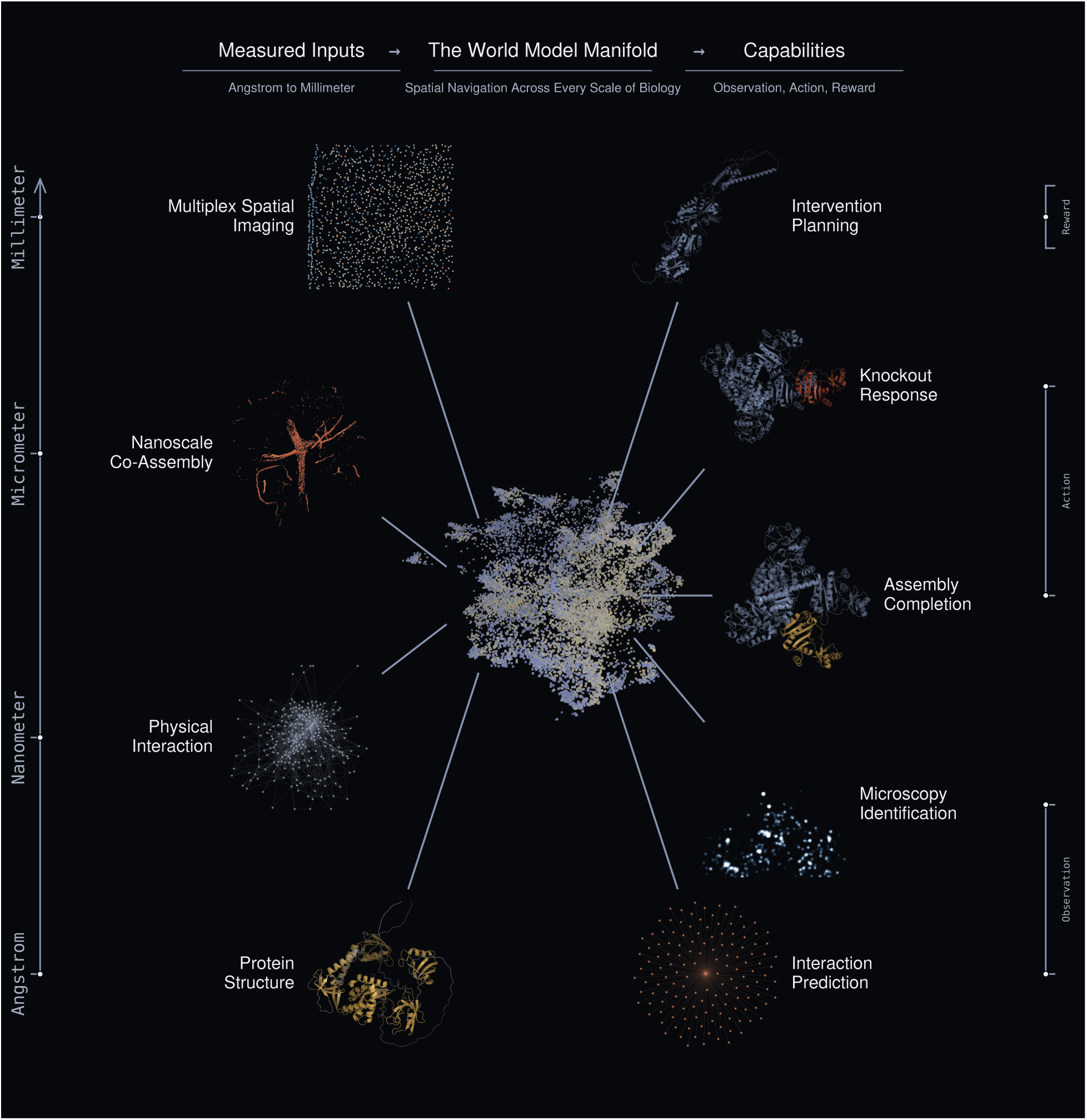
The HI-JEPA world model. Every panel is real output, not a schematic. Down the left are the measured channels the model learns from, ordered by spatial scale: a multiplex spatial field, a nanoscale co-assembly reconstruction, a physical-interaction graph and a protein structure. The center is the state itself, each point one protein embedding, colored by how much measurement supports it. Down the right are the operations read from that state, grouped by which world-model function they serve: observation, then action, then reward. Lines mark what feeds the state and what is read back out of it, and the rails in both margins mark where each panel sits on the scale ladder, with the arrow indicating that the ladder continues above the range reported here. What matters is that a tissue section and a protein backbone enter the same latent and the same range of scales comes back out.

**Figure 2.**
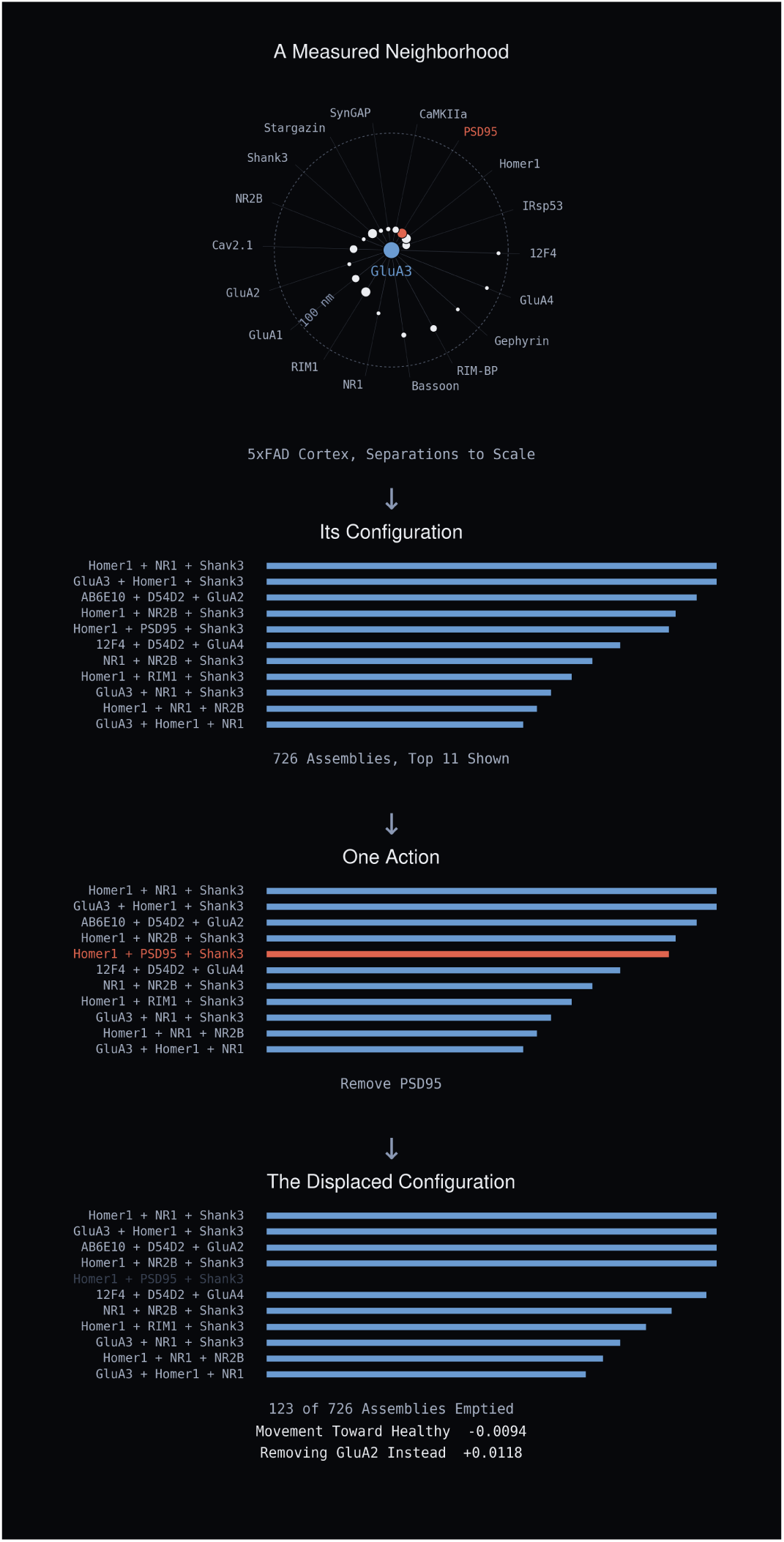
One measured neighborhood, end to end. A GluA3 cluster in 5xFAD somatosensory cortex has eighteen partners inside the 100 nm colocalization criterion, amyloid among them, drawn at their measured separations. The same neighborhood is then shown as the model holds it: abundance over a fixed assembly vocabulary, 726 of which are present here. Removing PSD95 empties the 123 assemblies that contain it and the remaining mass redistributes, which is what it means to displace a configuration. The model scores that displacement as movement away from the healthy state, while removing GluA2 from the same neighborhood moves toward it.

## 4. Data and Evaluation Protocol

### 4.1 Channels

ASCEND measures molecular proximity from nanoscale co-assembly to tissue architecture. Any modality that resolves molecular positions in intact tissue reduces to the same relation and enters the corpus through it, so the platform is tied to no single instrument. Expansion microscopy [23] supplies the highest-resolution proximity we hold. Multiplex spatial imaging carries proximity at coarser resolution, extending the corpus across more proteins and into human tissue. Reference channels cover physical interaction, co-function, co-disease, complex membership, deposited structure, perturbation response and compound bioactivity, each drawn from an established source and cited in Table 2. Every channel is partitioned by a strict train/held-out protein split before use.

**Table 1.**
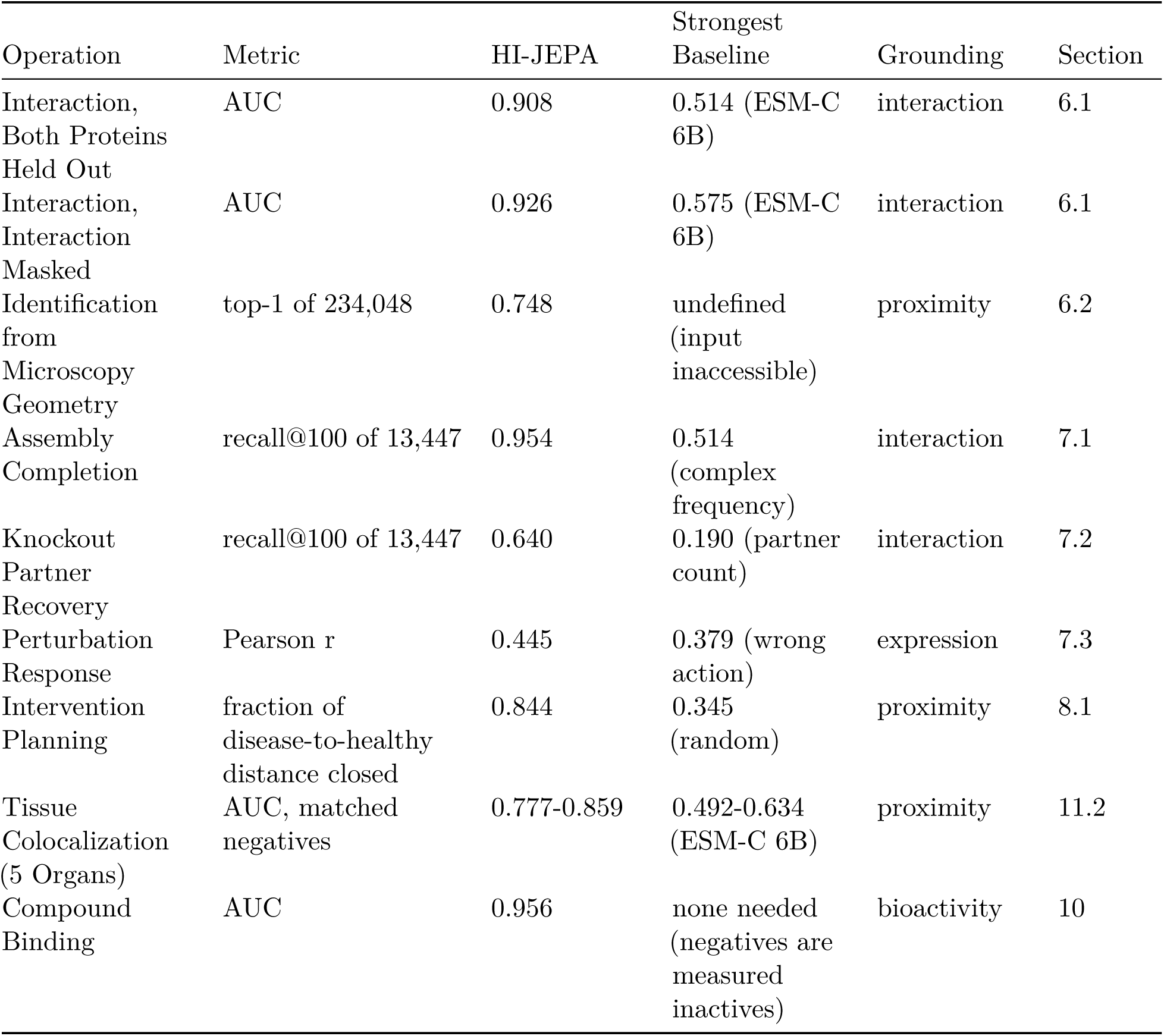
Summary of Principal Results.

**Table 2.** Channels and Sources.

| Relation | Source | Role |
| --- | --- | --- |
| Spatial Proximity (Primary) | Measured nanoscale co-assembly, ASCEND [23] | Navigation, assembly, grounding |
| Spatial Proximity (Breadth) | Multiplex spatial imaging | Cross-scale, tissue, coverage |
| Interaction | Experimental physical-interaction measurements [17] | Interaction |
| Perturbation | Held-out perturbation-response measurements [22] | Dynamics |
| Co-Function | Curated pathway annotation [18] | Function |
| Co-Disease | Curated disease-association references [19] | Disease |
| Complex Membership | Curated protein-complex reference [20] | Assembly completion |
| Structure | Deposited structural geometry [21] | Identification, cryptic |
| Drug-Target | Curated bioactivity measurements [24] | Binding |

ASCEND measures pairwise molecular proximity in intact tissue. The processed corpus spans five studies and 37 imaged regions, covering cultured neurons, isolated synapses, validation preparations and mouse somatosensory cortex in disease and control, over 164 proteins and roughly 1.2 million scored colocalization events.

One of those studies is public and serves as the worked example of how an acquisition becomes a measured relation: our reanalysis of the multiplexed expansion-revealing deposit of Kang, Schroeder, Lee et al. (2024), Harvard Dataverse doi:10.7910/DVN/JJBULY. Four 12-month-old mice, two 5xFAD and two wild-type, contribute nine diseased and eight control regions of primary somatosensory cortex, from which 161,678 nanoclusters were segmented and 40,831 colocalizations identified over 665 million candidate pairs. Clusters are segmented per channel by DBSCAN on a 99.9th-percentile intensity threshold with a voxel-aware neighborhood, and amyloid is retained only where three antibodies concur. Two clusters are colocalized when their voxel grids share or neighbor a cell, or their centroids lie within 100 nm pre-expansion, and each pair is scored against a density-matched label-shuffling null (n = 500) with Benjamini-Hochberg correction across the panel. Voxels are 162.5 x 162.5 x 250 nm post-expansion, 9.03 x 9.03 x 13.89 nm at the 18x expansion factor, and registration precision is 25 to 40 nm, so separations below that are reported as within the measurement limit rather than as exact distances. Measured proximity contributes 164 proteins and 0.04% of training edges, against 43.3% from multiplex spatial imaging and 56.7% from curated references.

Measured separations are reported as a distribution rather than as exact molecular distances. Median centroid separations pre-expansion are 22.4 nm for amyloid with GluA2, 82.9 nm for RIM1 with GluA2 in diseased tissue against 72.7 nm in control, and 89.7 nm for amyloid with RIM1, with the closest measured pair at 8.05 nm. Registration precision is 25 to 40 nm, so separations below that band lie within the measurement limit; only the distributions and the colocalization criterion are interpreted quantitatively.

### 4.2 Evaluation Protocol

All metrics are computed on held-out protein splits with 95% confidence intervals obtained by resampling. Metrics are named where reported: AUC over stated negatives, recall@k and top-1 against a given candidate pool, median rank in a list of stated length (lower better), Pearson r between predicted and observed change, cosine between predicted and measured vectors, and effective rank against the 384 dimensions available. Assembly completion, identification and binding use per-target splits. Identification uses a microscopy point cloud as the query, matched against the deposited structural proteome. Function and disease are scored against annotation sources independent of those used in training, and each value is checked for stability with the matching channel dropped. Inference within a trained checkpoint is deterministic, and capabilities reproduce across random initializations within measurement noise. Negative and boundary results are reported in full (Section 12).

**Baselines are floors, not chance.** Curated interactions and complexes concentrate on well-studied proteins, so a score using nothing but how often a candidate appears in them already clears chance by a wide margin. Wherever a readout ranks candidates or scores pairs, the reference point is a baseline built from the source data alone. Uniform chance is reported alongside only to indicate scale.

The split is a single random partition of the 13,447-protein vocabulary, fixed once and reused by every evaluation: 979 proteins held out, 12,468 in training, not stratified by family, pathway or degree. An edge is used only when both of its endpoints are training proteins, applied across every relation, and across all nine relations and 1,633,725 training edges the number that touch a held-out protein is zero. Where a pair appears in two relations it contributes two terms with separate weights; 13.8% of measured-proximity pairs are also curated, and the single dually-supported pair with both endpoints held out was never in training.

## 5. Training Objective

A predictive objective needs a paired term to prevent the representation from collapsing, and which term to use is decided by what the model is asked to do afterwards. We pair redundancy reduction with self-distillation (Barlow Twins [7], DINO [8]). The pairing was selected against the isotropic-Gaussian alternative on the capabilities of Table 3, so that table reports the comparison the choice was made on rather than an independent confirmation of it; the relative weighting of the two terms is part of the training procedure and is not described.

**Table 3.** Collapse-Prevention Objective. Both arms are trained separately for this comparison; neither is the shipped checkpoint. Everything matched but the objective. Higher is better except cross-scale rank.

| Capability | SIGReg | Barlow + DINO |
| --- | --- | --- |
| Spatial Navigation (Recall@100) | 0.280 | **0.771** |
| Interaction (AUC) | 0.659 | **0.911** |
| Function (Reactome Co-Pathway) | 0.611 | **0.731** |
| Disease (HPO Co-Phenotype) | **0.533** | 0.507 |
| Cross-Scale (Median Rank) | 8191 | **16** |
| Coverage of Newly Acquired Sources (Recall@100) | 0.017 | **0.072** |

The published isotropic-Gaussian alternative [6] is provably optimal for a linear probe and flattens the density toward uniform. Our readouts are nearest-neighbor retrieval over a relation graph and need the clustered geometry that flattening removes, so it costs nearly every retrieval metric and sends cross-scale rank from 16 to effectively random (Table 3). The single exception is disease co-phenotype, where the flattened representation scores 0.533 against 0.507, consistent with a linear-probe-shaped readout being the one case flattening helps.

A contrastive control trained on identical data and relations isolates the effect of the objective (Table 4). HI-JEPA leads on every capability, by a wide margin on navigation and interaction and narrowly on function.

**Table 4.** Capability Comparison. Shipped checkpoint. Higher is better except cross-scale rank.

| Capability | Contrastive Baseline | HI-JEPA |
| --- | --- | --- |
| Spatial Navigation (Recall@100) | 0.748 | **0.847** |
| Interaction (AUC) | 0.859 | **0.910** |
| Function (Reactome Co-Pathway) | 0.875 | **0.881** |
| Cross-Scale (Median Rank) | 21 | **14** |

Dimensional collapse is the principal failure mode of non-contrastive self-supervision. The learned representation holds 2.5 times the effective rank of the contrastive control on navigation and 3.9 times on interaction (each modality is encoded to a 768-dimensional latent and the read-out heads project to 384, so an effective rank of 121.6 occupies roughly a third of the available dimensions).

The data-efficiency gap follows from the objective. InfoNCE learns a manifold by separating measured positives from generated negatives, so every recorded proximity determines the 2,000 negative pairs drawn against it. The 2,000x figure and the 2,000 negatives per positive are therefore the same quantity seen from two sides, not two independent findings. HI-JEPA predicts a measured neighbor’s embedding from the query protein and scores only that positive. Trained to comparable interaction AUC (0.92 against 0.90), the contrastive control processes roughly 2,000 times more pair comparisons. On a biological graph the generated negatives are frequently real interactions that happen to be unlabeled, so the contrastive objective penalizes pairs that genuinely interact.

**Figure 3.**
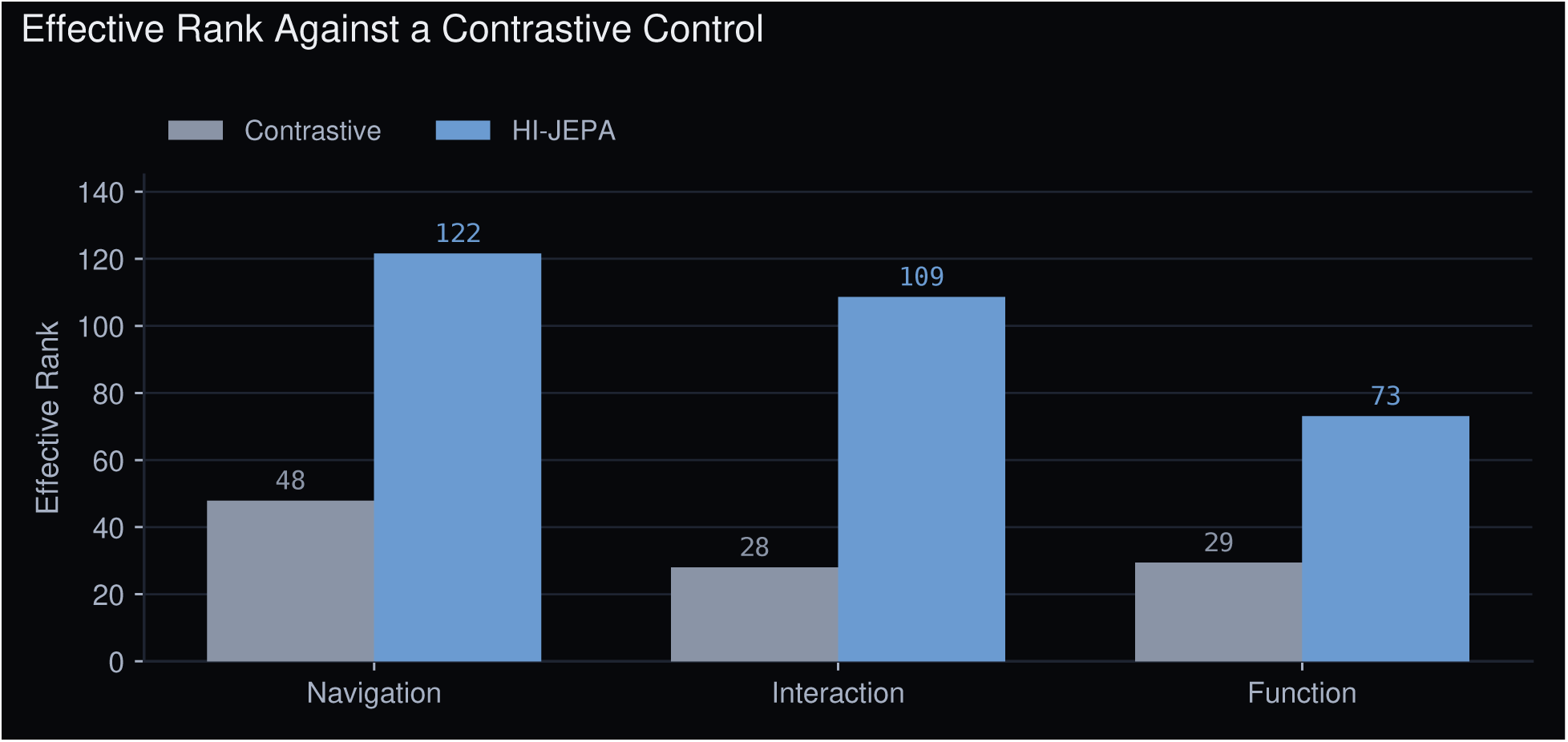
Effective rank against a contrastive control trained on identical data and relations, counting how many latent dimensions carry independent signal. Higher is better. Capability scores are Table 4.

## 6. Identity from Measurement

HI-JEPA places a real measurement into its state in two cases a sequence model cannot handle: interactions between proteins whose sequences are uninformative, and raw microscopy, which the sequence channel cannot read at all.

### 6.1 Interaction Prediction for Sequence-Dissimilar Pairs

We evaluate interaction prediction where sequence methods are least reliable, on high-confidence physical interactions between proteins whose sequences are dissimilar, and compare against ESM-C [10] at 600M and 6B parameters, varying how dissimilar the pairs must be. HI-JEPA holds 38.9M parameters, so the comparison runs against sequence models roughly fifteen and one hundred and fifty times its size. The result is a curve rather than a single cut.

Two protocols are reported (Table 5). The first withholds both proteins of a test pair from training, so the model has never seen either in any context. The second masks only the interaction between them, leaving each protein visible on its other interactions. The masked-interaction split is the conventional one and supplies a much larger test set; withholding both proteins is the harder test.

**Table 5.**
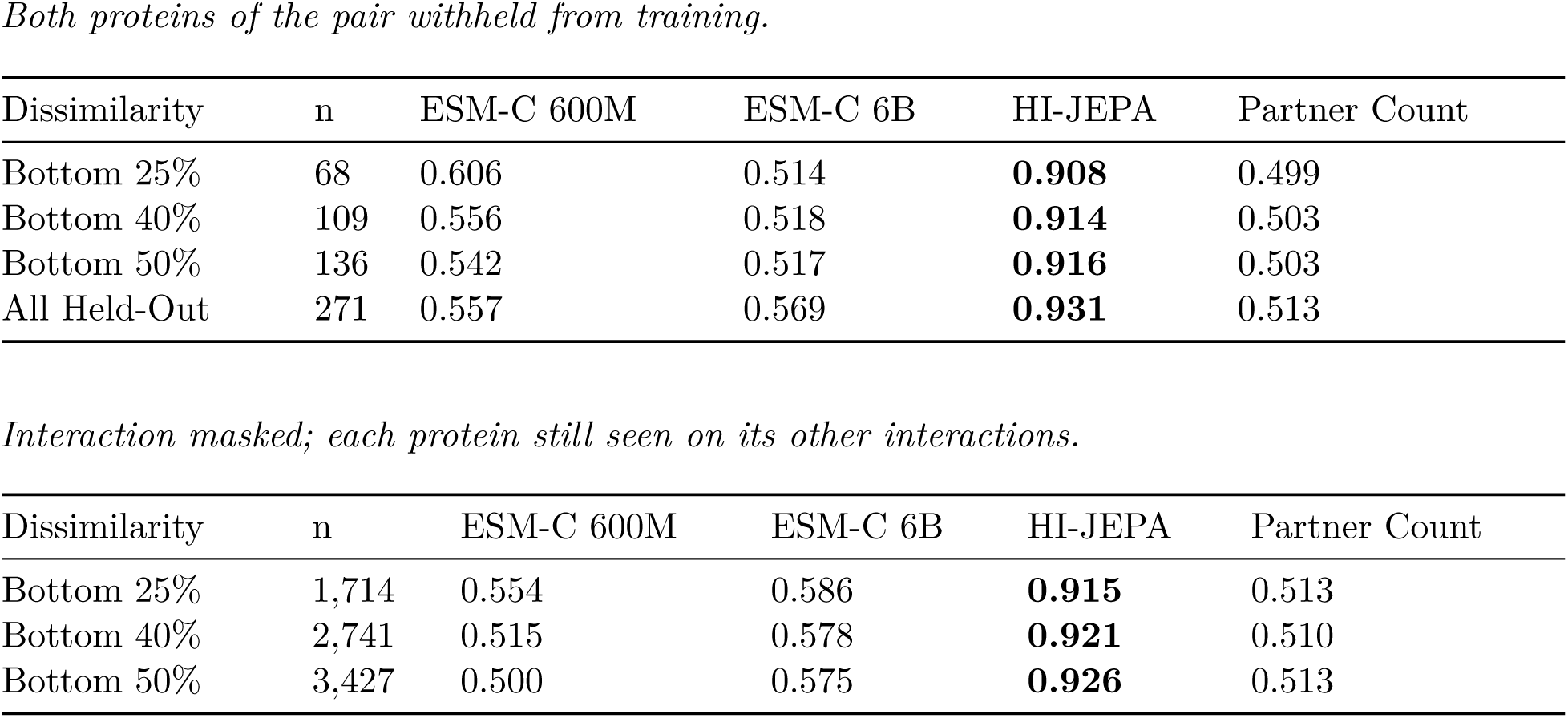
Interaction AUC on Sequence-Dissimilar Interacting Pairs. Shipped checkpoint. Positives are high-confidence physical interactions. Negatives are non-interacting pairs drawn at the same dissimilarity threshold and matched to the positives on partner count. Dissimilarity is the bottom percentile of pairwise sequence-embedding cosine. Values are the mean over twelve independent negative draws; the spread across draws is at most 0.030. The last column scores a pair using nothing but its two partner counts.

| Dissimilarity | n | ESM-C 600M | ESM-C 6B | HI-JEPA | Partner Count |
| --- | --- | --- | --- | --- | --- |
| Bottom 25% | 68 | 0.606 | 0.514 | **0.908** | 0.499 |
| Bottom 40% | 109 | 0.556 | 0.518 | **0.914** | 0.503 |
| Bottom 50% | 136 | 0.542 | 0.517 | **0.916** | 0.503 |
| All Held-Out | 271 | 0.557 | 0.569 | **0.931** | 0.513 |

*Interaction masked; each protein still seen on its other interactions.*
| Dissimilarity | n | ESM-C 600M | ESM-C 6B | HI-JEPA | Partner Count |
| --- | --- | --- | --- | --- | --- |
| Bottom 25% | 1,714 | 0.554 | 0.586 | **0.915** | 0.513 |
| Bottom 40% | 2,741 | 0.515 | 0.578 | **0.921** | 0.510 |
| Bottom 50% | 3,427 | 0.500 | 0.575 | **0.926** | 0.513 |

A protein with many known partners appears in many positive pairs, so negatives drawn without regard to partner count let a score built from the two counts alone reach 0.619. Matching the negatives on partner count sends that score to 0.499. The same correction takes HI-JEPA from 0.917 to 0.908 and ESM-C 6B from 0.607 to 0.514: most of what the sequence baseline appeared to know about these pairs was partner count.

The two model scales separate the question from capacity. Where one partner is still seen on its other interactions, the ten-fold larger model leads the smaller by 0.031 to 0.075. Where both partners are withheld, that ordering reverses, the 600M model leads the 6B by as much as 0.092, with both near the partner-count floor. Ten times the parameters gains nothing once the familiar partner is removed. What is missing from the sequence is not capacity.

**Figure 4.**
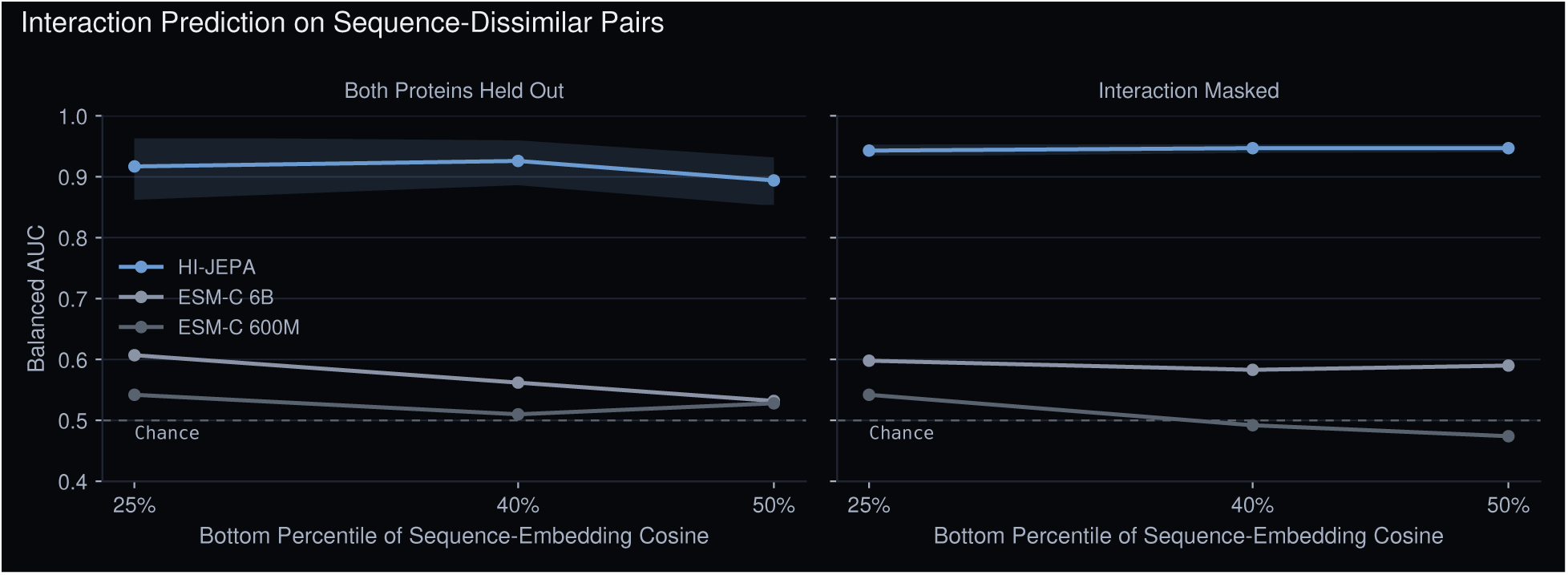
Interaction AUC on sequence-dissimilar pairs against the dissimilarity threshold, under both protocols. Both scales of ESM-C remain near the partner-count floor while HI-JEPA attains 0.89–0.95 with non-overlapping confidence intervals at every threshold, including the 3,427-pair masked-interaction split.

#### 6.1.1 Removing the Protein Family

Holding a protein out does not by itself remove its family: a held-out kinase is still scored against a training set full of kinases, which leaves family recall as an explanation. Clustering the vocabulary at 30% sequence identity separates the two. Of 979 held-out proteins, 375 have no relative anywhere in training (clustered over the full 13,447-protein vocabulary, which yields 7,069 families at 30% identity). On pairs where neither partner has a relative, HI-JEPA reaches 0.947 against 0.906 where one was available, and the same ordering holds at every dissimilarity threshold. Removing the family raises the result rather than lowering it. ESM-C moves the other way and falls to 0.414 on the hardest cell, which is what a model relying on family recall would do.

#### 6.1.2 Against a Co-Folding Model

Sequence similarity is also what a co-folding model has to work with. ESMFold2 predicts the complex itself rather than reading a representation, and its authors report it exceeding established methods on biomolecular complex prediction [10]. On these pairs it reaches 0.538, with an interval spanning chance.

*Co-folding, both proteins held out*.

| Model | n (Pos / Neg) | AUC |
| --- | --- | --- |
| ESMFold2 (Interface Confidence)* | 49 / 48 | 0.538 [0.420, 0.654] |
| HI-JEPA on the Identical Pairs* | 49 / 48 | **0.900 [0.828, 0.961]** |

\* Subsample, single negative draw. fp32 co-folding exceeds a single 141 GB GPU above roughly 2,900 combined residues, so ESMFold2 is scored on the 97 pairs of the strict protocol whose combined length fits, and HI-JEPA is rescored on exactly those pairs. Folding is expensive enough that these two rows use one draw with resampled intervals rather than the twelve draws behind every other row; the ESMFold2 interval spans chance either way.

Every positive here is an experimentally documented physical interaction. The mean interface confidence across them is 0.144, with none above 0.8 and one above 0.5, so the model returns no usable complex for interactions that are known to form.

Sequence methods extrapolate from similarity, and a target is worth pursuing precisely when it resembles nothing well characterized, so the prediction is weakest where the value is highest. Measured proximity is an observation rather than an extrapolation and carries no similarity requirement.

### 6.2 Identification from Microscopy Geometry

The model names a protein from the bare geometry of a microscopy point cloud, with no sequence, no label and no antibody raised against it. Sequence and co-folding models require an amino-acid sequence and cannot accept a point cloud, which settles that comparison.

A query is the microscopy point cloud of one held-out protein, matched against the deposited structural proteome. Identities in the measured corpus are assigned by antibody, so no geometry-derived label enters training and the evaluation here is scored against deposited structures, a source disjoint from the panel that defines the corpus. Recognition spans a separate physical instance of the same protein, at a different construct, conformation and resolution. A field of view contains many copies of each protein, and accuracy is also reported with those copies pooled.

**Table 6.**
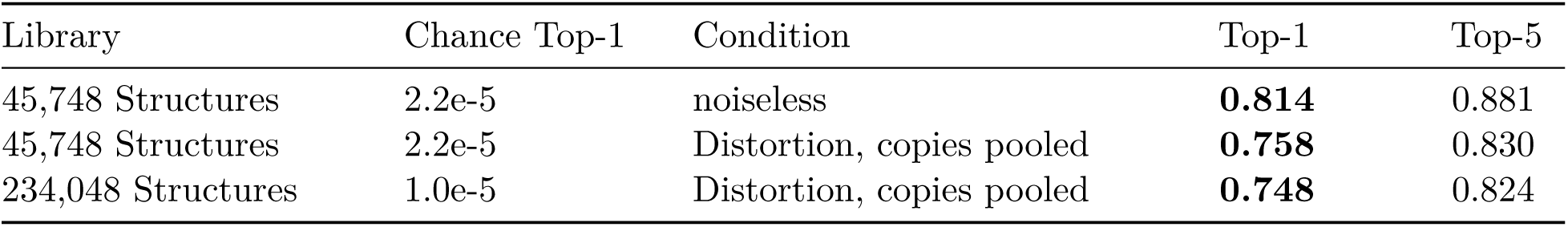
Antibody-Free Identification against the Structural Proteome.

| Library | Chance Top-1 | Condition | Top-1 | Top-5 |
| --- | --- | --- | --- | --- |
| 45,748 Structures | 2.2e-5 | noiseless | **0.814** | 0.881 |
| 45,748 Structures | 2.2e-5 | Distortion, copies pooled | **0.758** | 0.830 |
| 234,048 Structures | 1.0e-5 | Distortion, copies pooled | **0.748** | 0.824 |

Pooling the multiple copies a real field of view provides recovers most of the noiseless accuracy under distortion, because the noise averages as copies accumulate. Flooding the library to 234,048 deposited structures (adding 52,000 complexes and unmapped structures as pure distractors) costs a single point of top-1.

**Figure 5.**
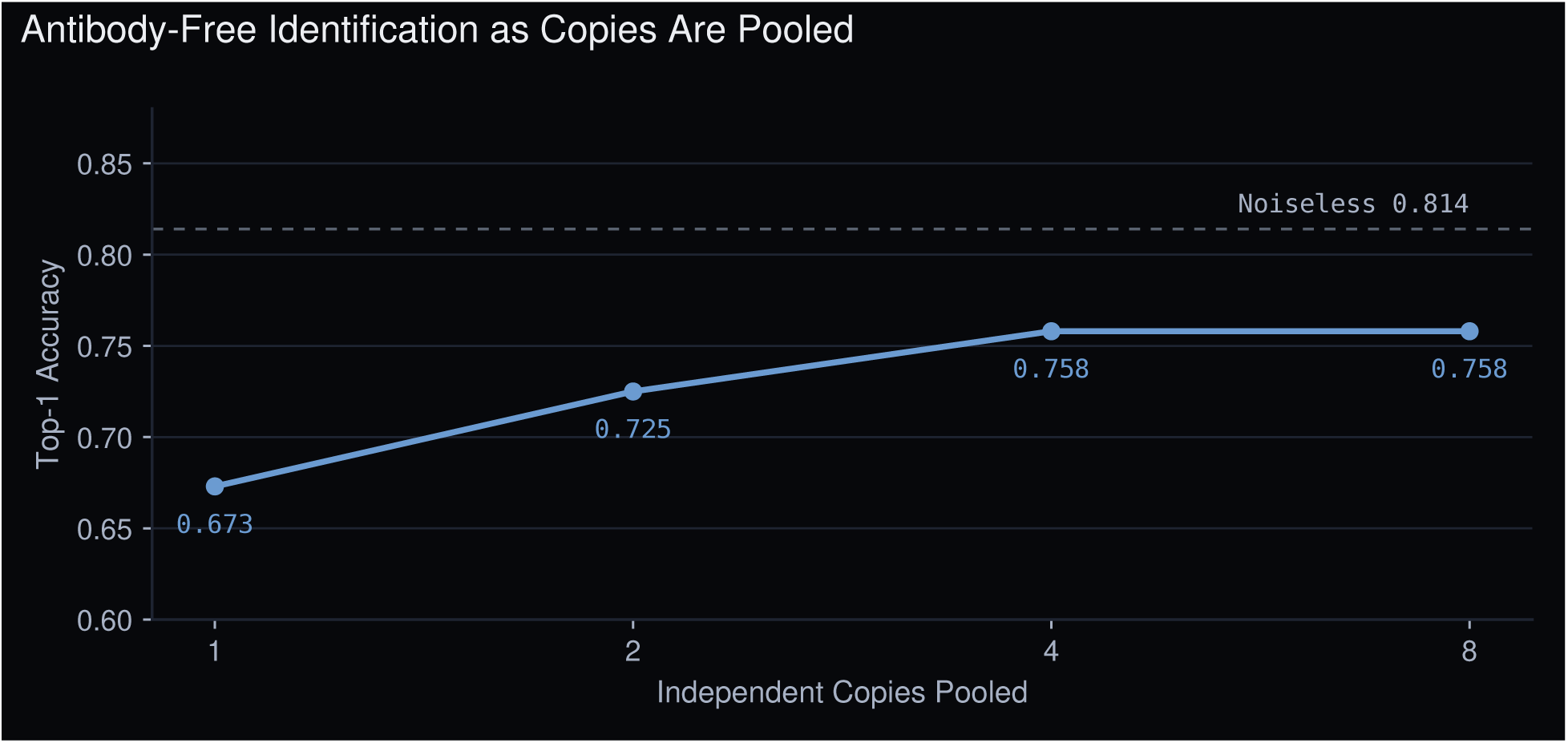
Antibody-free identification against 45,748 deposited structures under realistic measurement distortion. Top-1 accuracy rises as independent copies are pooled, recovering most of the noiseless performance at the realistic operating point; chance is 2.2e-5. Flooding the library to 234,048 structures costs a single point.

## 7. Acting on the State

Each result below applies an action to a configuration and reads the consequence. A complex has a member taken out; a protein is removed from its neighborhood; a perturbation is carried forward under a learned transition. Completion and knockout are the same operation in opposite directions, one asks what fills a gap, the other what a gap disturbs.

### 7.1 Assembly Completion

Given a partial complex, the model ranks candidate missing members. On a literature-curated complex reference (EBI Complex Portal [20]) with zero-shot held-out targets (n = 325), a truly held-out member lands in the top 100 of 13,447 candidates at recall 0.954 [0.929, 0.975], and in the top 10 at 0.492.

Curated complexes concentrate on well-studied proteins, so the floor sits well above chance: ranking candidates by how many complexes each already appears in reaches 0.514 at 100 and 0.246 at 10. HI-JEPA is 1.9 times that floor. Removing the pathway-annotation channel leaves it at 0.917. On the identical task ESM-C 6B reaches 0.141, below the floor that counting complex memberships already clears.

### 7.2 Knockout Response

The model names the partners disrupted by removing a protein, ranking 13,447 candidates by latent proximity to the knocked-out protein. On 62 targets with the target and its partners both held out, this recovers the experimentally observed partners at recall@100 0.640 [0.55, 0.72].

The action-conditioning control is the sharpest in the paper. Ranking against a different protein’s embedding, with the targets, the ranking rule and the denominators all unchanged, collapses recovery to 0.028 [0.00, 0.07], and the intervals do not overlap. Asked about the wrong protein, the model does worse than a sequence embedding. A ranking built from partner counts alone returns the same ordering whichever protein was removed (it is blind to the action by construction) and recovers 0.190. Both scales of ESM-C fall below that at 0.119 and 0.047. At the top ten, partner counts recover nothing while HI-JEPA reaches 0.132.

**Table 7.** Knockout Partner Recovery. Shipped checkpoint. Recall of experimentally observed physical partners among the top-ranked candidates of 13,447; 62 held-out knockout targets; 95% confidence intervals. The fourth row repeats the model’s own ranking against a different target’s embedding, changing only the action. The fifth orders candidates by partner count and ignores the knocked-out protein, measuring how much recall comes from well-studied proteins being over-represented among observed partners.

| Model | Recall@10 | Recall@50 | Recall@100 |
| --- | --- | --- | --- |
| HI-JEPA | **0.132** [0.09, 0.18] | **0.478** [0.41, 0.55] | **0.640** [0.55, 0.72] |
| ESM-C 6B | 0.026 [0.00, 0.05] | 0.077 [0.04, 0.12] | 0.119 [0.07, 0.17] |
| ESM-C 600M | 0.022 [0.01, 0.04] | 0.039 [0.01, 0.07] | 0.047 [0.02, 0.08] |
| Another Target’s<br>Action | 0.012 [0.00, 0.03] | 0.027 [0.00, 0.07] | 0.028 [0.00, 0.07] |
| Partner Count Only | 0.000 | 0.133 [0.09, 0.19] | 0.190 [0.13, 0.26] |
| Chance | 0.0007 | 0.0037 | 0.0074 |

### 7.3 Action-Conditioned Dynamics

A transition takes a state and an action and returns the state that follows, *z*^′^ *= F(z, a)*. This section tests the action half of that operator in a second measurement space. The state here is a population of cells rather than a configuration of proteins, and the prediction is how far each measured gene’s expression moves; what the world model supplies is the action, which is the embedding of the silenced gene. Because the question is what an action produces rather than when, the transition carries no time index.

Testing the same operator against a different observation space is deliberate. If action conditioning only ever appeared where the model was trained, it would be difficult to separate from retrieval. The comparison at the end of this section is the point: the same substitution costs far more in the space the measurement is taken in.

A transition could score well while ignoring what it is told, by learning the average response to a perturbation and returning it regardless, so each input is replaced with a wrong one in turn. On 2,749 held-out combinations of a silenced gene and a cell line [22] the transition reaches 0.445 (Pearson correlation between predicted and observed expression change, computed per held-out record and bootstrapped over records). Substituting another gene’s action drops it to 0.379; applying the perturbation in the wrong cell line drops it to 0.330. Neither interval overlaps the first.

Every silenced gene produces some common consequence, and a model can score well on that shared component without knowing which gene was hit. Removing the mean response leaves only what is specific to the perturbation. On that residual the transition reaches 0.361 [0.345, 0.376] against 0.200 [0.185, 0.217] for another gene’s action, so the margin attributable to knowing the action more than doubles once the shared component can no longer carry the score.

A learned transition also predicts the configuration itself. On 17 measured regions with one region held out in turn (1,574 assemblies over 23 markers), it reproduces the held-out configuration at cosine 0.947 (cosine similarity between the predicted and the measured assembly-abundance vectors), against 0.933 for a mean-configuration baseline and 0.883 for a mass-action one, so a region’s configuration is predictable from the others and not only from average composition.

The same substitution is far sharper in the space the measurement is taken in. Recovering the partners a knockout disrupts asks the same question of a configuration rather than of expression, and there the identical control takes the model from 0.640 to 0.028 (Section 7.2). Substituting the action costs 0.066 in expression and 0.612 in measured proximity: the conditioning is strongest where the measurement is.

**Figure 6.**
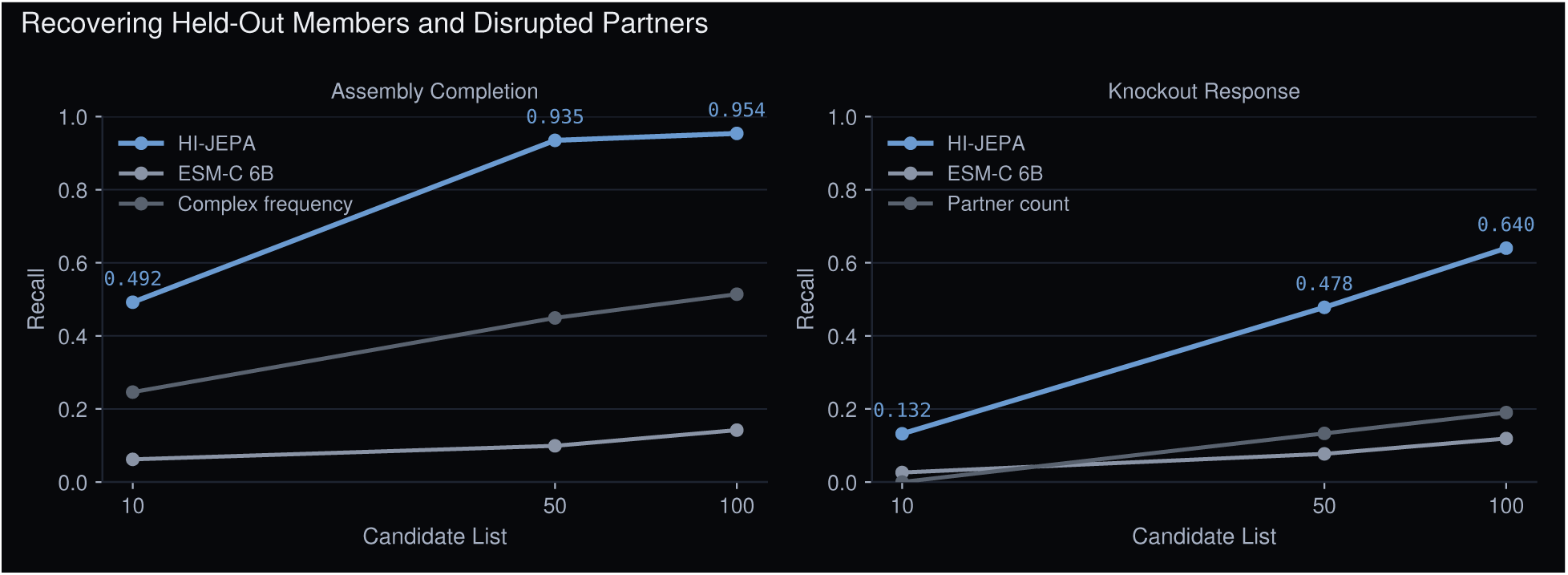
Acting on the state. Completing a partial assembly and naming the partners a knockout disrupts, both ranking the same 13,447 candidates under the same rule, against a sequence model and against a ranking that ignores which protein was involved.

## 8. Planning against a Measured Goal

### 8.1 Intervention Planning

Planning searches over single-protein interventions for the sequence that moves a diseased configuration toward a healthy one, scored along an axis fit from paired 5xFAD [25] and wild-type tissue. Searching over a horizon closes 0.844 of the distance to the healthy state (95% CI [0.696, 0.984]), against 0.345 for random interventions and 0.019 for a single action, all three scored by the identical rule. The intervention model is definitional rather than fitted: removing a protein empties the assemblies built from it and the remaining mass renormalizes, since an assembly containing a removed species cannot be observed. The learned transition over these same configurations is reported in Section 7.3.

What the plan selects is specific to that axis. It disrupts all three A*β*–AMPA subunits (GluA2, GluA3, GluA4), which the published literature independently identifies as the disease’s principal mediators (Section 8.3). Permuting the 5xFAD and wild-type labels across the 17 regions and refitting the axis gives the rate at which a meaningless direction lands on all three by chance: 4% of 1,000 permutations (p = 0.041). The regions come from four animals and are not independent within an animal, which makes the permutation one over regions rather than animals (S1 contributes five regions and S2 four, both 5xFAD; S3 and S4 four each, both wild-type. The axis is fit on all seventeen regions, including those it then scores, which is what the permutation test controls for). The same permuted axes still close 0.740 of the distance on average, so the distance on its own does not carry the result, the subunit selection does.

### 8.2 Human Disease Translation

Planning scores an action by displacement along a measured proximity axis between a diseased and a healthy configuration. That goal is meaningful only if measured proximity carries disease state in human tissue, and not only in the tissue the model was trained on.

**Figure 7.**
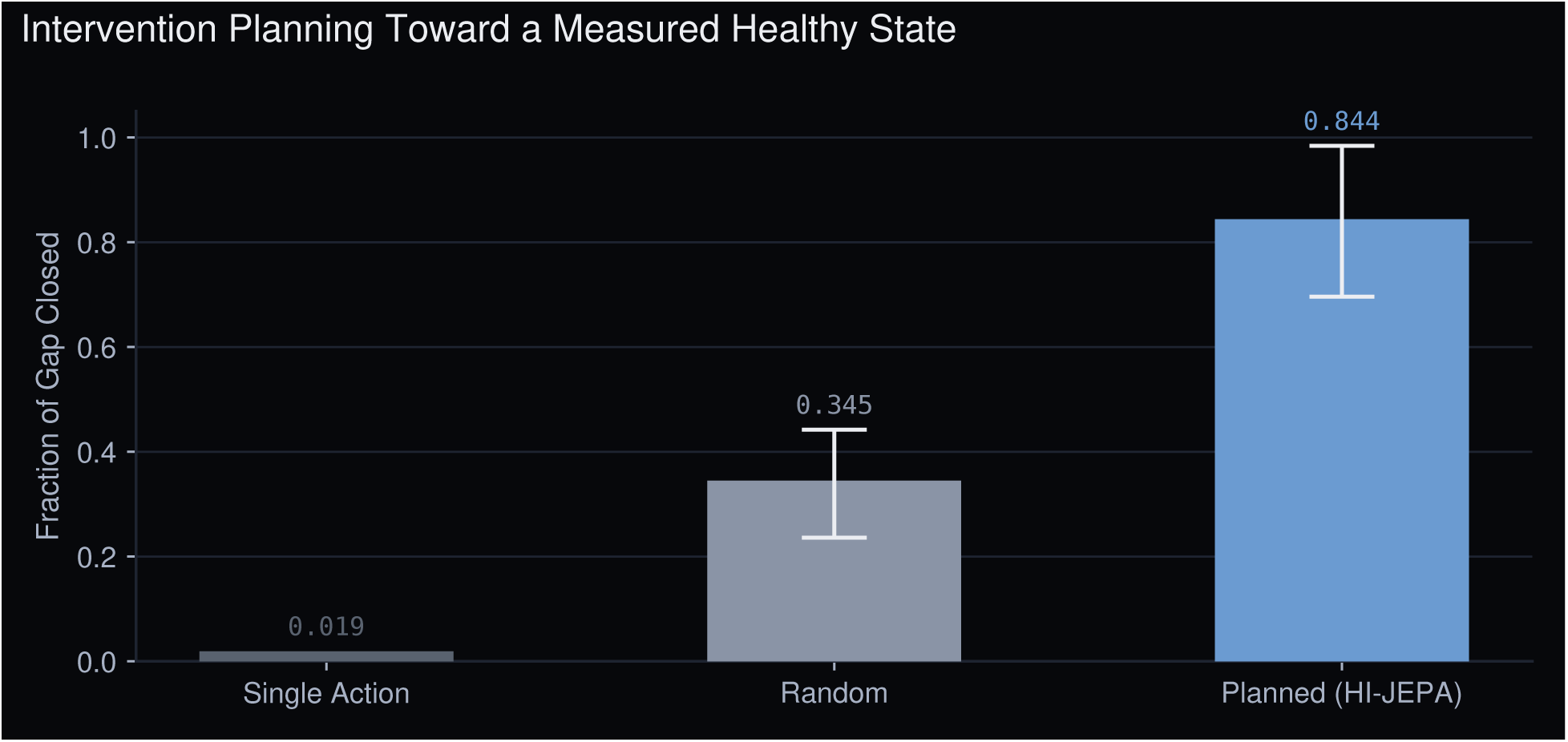
Intervention planning. Displacement toward the healthy state for a single action, random interventions, and the planned HI-JEPA trajectory; error bars are 95% confidence intervals.

Proximity is read from human tissue as it is read for training: by scoring how much more two species colocalize than that sample’s own geometry produces by chance, and correlating that score against donor severity across a cohort. Any spatial modality that resolves proximity admits the same read. On held-out human cohorts the axis holds with no further training, and each pair carries a direction fixed before the test.

· In spatial transcriptomics of Alzheimer’s brain, TREM2–APOE colocalization separates the five most severely affected donors from the five least by a z-score difference of 3.6 [1.1, 5.9], recovering the disease-associated-microglia signature; across all ten donors it tracks severity continuously at r = 0.591.
· In a second cohort of 27 donors, astrocyte colocalization with upper-layer neurons separates severe from mild by a z-score difference of 14.9 [6.2, 23.9] at two-sided p = 0.002, recovering reactive astrogliosis.
· In a seven-donor liver cohort, stellate cells and fibrotic hepatocytes separate diseased tissue from healthy by a z-score difference of 5.3 [2.2, 8.9].
· The Alzheimer’s result replicates in an independent 32-donor cohort on a different platform.

Every interval excludes zero. The cohorts are small, and on the smallest a permutation test resolves no finer than 0.08 two-sided, so what carries the claim is four independent datasets across three modalities and three organs agreeing in the direction each specified in advance. Cross-condition contrasts frequently do not reach significance and are reported in Section 12.

The quantity the planner optimizes therefore carries disease state in humans.

### 8.3 Corroboration of Predicted Dynamics

From measured nanoscale geometry alone, the model ranks disruption of the A*β*–AMPA coassemblies (GluA2, GluA3, GluA4) as therapeutic and predicts that disruption of the PSD-95 scaffold worsens the configuration. Both predictions agree in sign with the experimental literature, and the subunits selected are the ones the literature implicates. The planner ranks interventions by displacement along the measured axis, which is a different quantity from per-pair amyloid enrichment, so the two orderings are not expected to match subunit for subunit; what agrees is that AMPA subunits are selected and that PSD-95 goes the other way. A*β* induces synaptic depression by removing AMPA receptors, and inhibiting that endocytosis rescues the deficit; GluA3-containing receptors are required for A*β*-driven synaptic and memory loss, and GluA3-null neurons are resistant [1, 2]. PSD-95 protects synapses from A*β*, and its reduction is associated with postsynaptic degeneration [3].

### 8.4 Target Nomination from Anomalies

A target list is ordinarily downstream of a hypothesis, which is exactly what a poorly understood disease does not have. ASCEND resolves which proteins occupy the same nanoscale neighborhood in diseased and healthy tissue from one experiment, which makes the disease’s rearrangement directly visible. Target nomination therefore reduces to a ranking.

We measure 1,574 assemblies, each a set of proteins seen together in one neighborhood, and rank them by how far their abundance in 5xFAD departs from wild type. The ranking uses no reward, no plan, no disease label, and no indication that the tissue is Alzheimer’s or that amyloid is relevant.

Nine of the top ten are amyloid-*β* bound to AMPA receptor subunits, the assemblies the literature identifies as the mediators of the disease (Section 8.3). Of the 1,574, 206 carry that signature, so ten drawn at random return one or two against nine for this ranking. Abundance is the natural baseline, since a common assembly moves further in absolute terms; ranked by abundance the same depth returns 0.300. Enrichment runs 6.9-fold at ten, 3.4 at twenty, 2.3 at thirty, and returns to the base rate by one hundred.

Three independent routes through the system reach GluA2 and GluA4: this ranking from the measured configuration alone, the planner of Section 8.1 searching over the same configurations, and the published literature from experiments the model never saw. The first two share the same underlying measurement, so they are not fully independent of each other; the third is.

Alzheimer’s serves as the test case because its mediators are independently established. Recovering them from tissue alone is what licenses applying the same procedure where they are not yet known.

## 9. Ablating Measured Proximity

A good representation that had simply seen more data could account for every capability above. Removing the measurement distinguishes that explanation from grounding. Each measured channel is ablated at training time, and we then score two capabilities that a merely good representation would not separate: cross-scale partner recovery and held-out within-scale dynamics.

· Removing measured proximity degrades cross-scale recovery (median rank 14 → 68) while navigation holds (0.847 → 0.853) and within-scale dynamics hold (median rank 1,164 → 1,268 of 13,447 candidate interventions).
· Removing perturbation degrades within-scale dynamics (median rank 1,164 → 2,403) while cross-scale recovery holds at 14.

Each channel produces one capability and leaves the other alone. The perturbation arm is the positive control: the assay detects a dynamics hit when one exists, so the flat dynamics result under proximity ablation is a real negative rather than an insensitive test.

Measured proximity is 0.04% of training edges, but it is the only relation whose endpoints sit at different scales, so a cross-scale operation has no other source to learn from. Masking the channel at inference leaves cross-scale recovery unchanged, so the model is not retrieving those edges at query time. The test also requires a measured proximity channel to remove, which a manifold assembled from foundation-model similarity or a knowledge graph does not contain.

The channel carrying the cross-scale operation is the one we acquire ourselves. Extending the corpus, rather than the architecture, is therefore the more productive direction.

**Figure 8.**
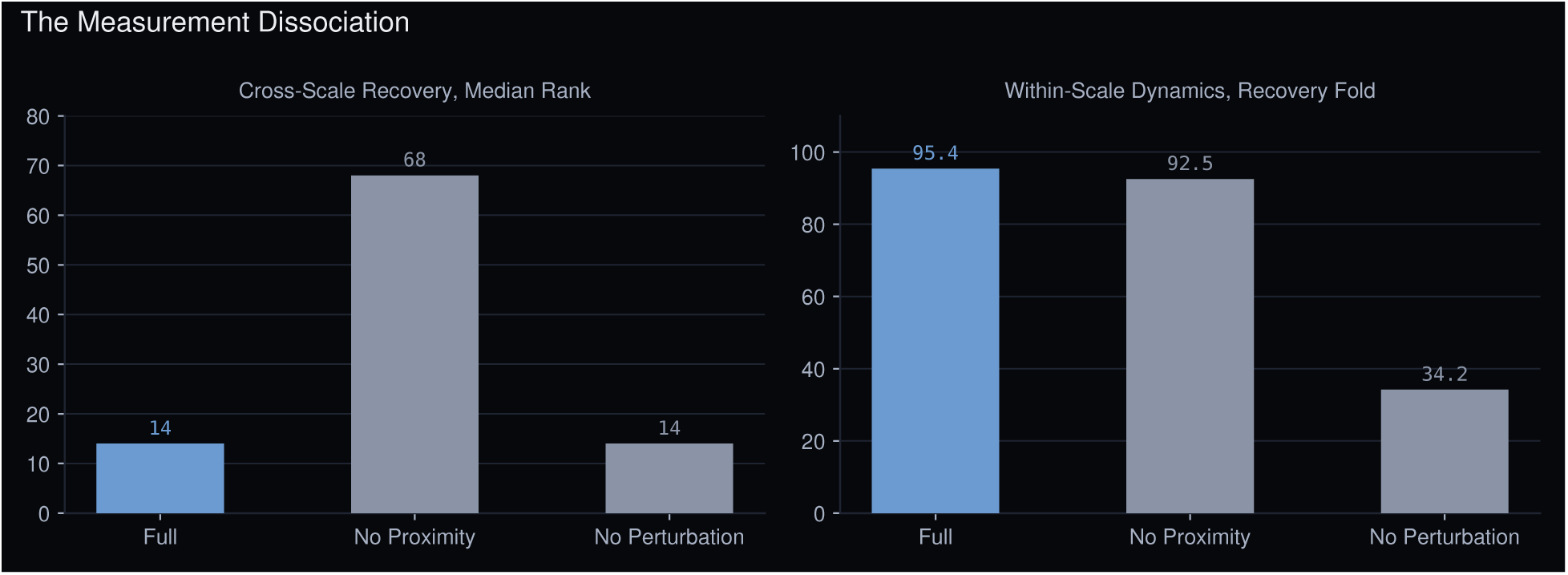
The measurement dissociation. Removing measured proximity degrades cross-scale partner recovery (median rank, lower better) while navigation and dynamics hold; removing perturbation degrades dynamics while cross-scale holds.

## 10. Performance across Biological Scales

One latent carries every scale of the biological hierarchy; that is what this paper means by *hierarchical*. The alternative is a model per scale and a pipeline to join them, where every crossing needs its own mapping, every mapping needs its own training data, and errors compound along the chain. Here a crossing is a distance in one space, computed the same way at every scale. Every number below is computed on held-out data and appears with its provenance in Table 1.

· **Molecular**, Compound–protein binding, AUC 0.956, compound held out (held-out-compound split over 13,811 compound-protein pairs, positives at pchembl >= 6 and negatives measured inactive at pchembl < 5 rather than sampled decoys; against a popularity-debiased shortlist the true target’s median rank is 6 of 13,447).
· **Protein**, Interaction 0.91–0.93 between proteins never seen, where a sequence embedding sits at its partner-count floor (§6.1). A microscopy point cloud maps to identity against 234,048 structures at top-1 0.748 (§6.2).
· **Complex**, A held-out member recovered in the top 100 at 0.954, against 0.514 from complex frequency alone (§7.1); triple co-assembly does not follow a mass-action composition law, since assembly abundance scales with the product of its constituents at an exponent of 0.64 [0.60, 0.68] rather than 1.0, and abundance alone leaves 46% of the variance unexplained.
· **Cellular**, Response to silencing a gene predicted at 0.445 across 2,749 held-out gene and cell-line pairs, against 0.379 when the action is replaced (§7.3).
· **Tissue**, Colocalization AUC 0.777–0.859 across five organs under partner-matched negatives, against 0.49–0.62 from partner counts alone (§11.2).

## 11. Comparison with Sequence Models

Every operation above that involves measured proximity or a configuration can also be posed to a sequence model, by ranking the same candidates on sequence-embedding similarity over identical held-out data. Wherever the two compare, the strongest sequence models sit near their floor. Microscopy-native identification is the exception: the comparison is undefined because the input is a point cloud a sequence model cannot accept.

These operations need measured proximity, and a sequence model has no channel that carries it. What is tested is therefore how far a sequence representation generalizes, not how well it performs the task it was trained for. Stating that plainly matters, since the comparison would otherwise read as unfair.

### 11.1 Supervised Probes

The sequence baselines above read out by embedding similarity, which is unsupervised, while HI-JEPA’s readouts are trained. We therefore match the supervision: freeze ESM-C 6B, train a probe on the same labels over the same held-out split, and evaluate on identical pairs (Table 8).

**Table 8.**
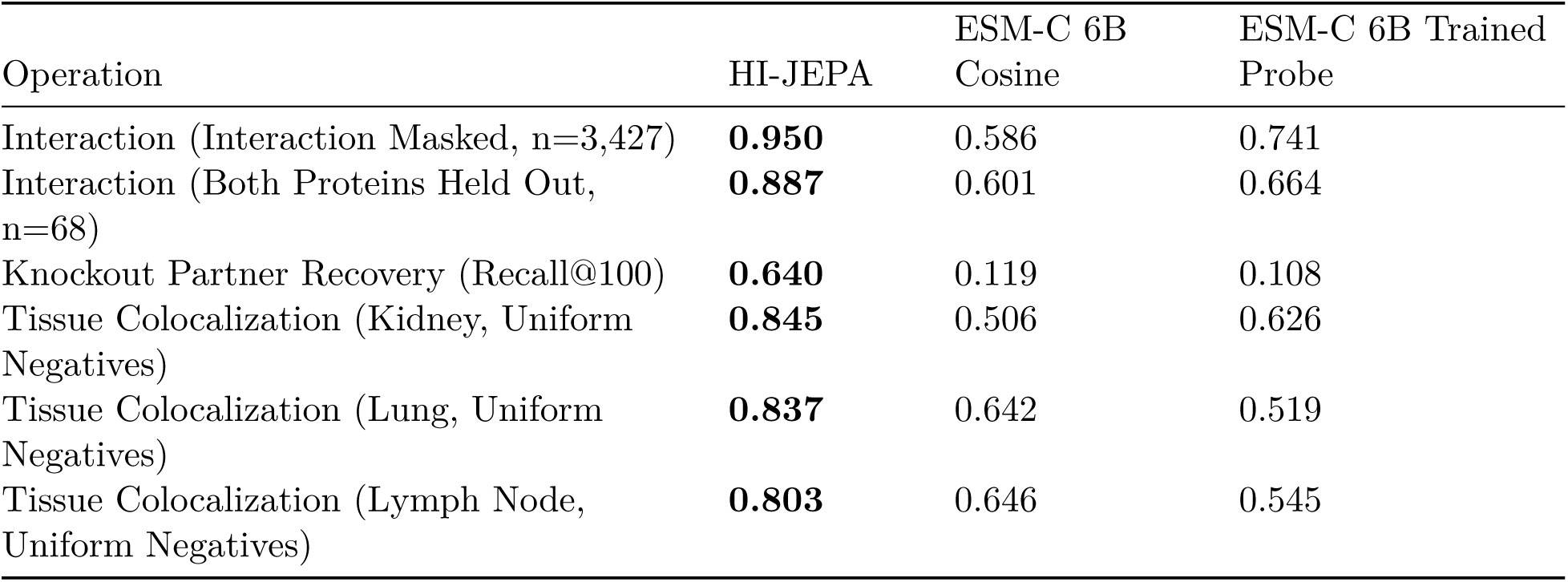
ESM-C 6B with and without a Supervised Probe. Shipped checkpoint. Both scored on identical held-out pairs, by cosine similarity and by a probe trained on the same labels. HI-JEPA on the same pairs for reference. Negatives here are drawn uniformly rather than matched on partner count, so all three columns sit above their Table 5 values and the comparison holds within this table only.

| Operation | HI-JEPA | ESM-C 6B<br>Cosine | ESM-C 6B Trained<br>Probe |
| --- | --- | --- | --- |
| Interaction (Interaction Masked, n=3,427) | **0.950** | 0.586 | 0.741 |
| Interaction (Both Proteins Held Out, n=68) | **0.887** | 0.601 | 0.664 |
| Knockout Partner Recovery (Recall@100) | **0.640** | 0.119 | 0.108 |
| Tissue Colocalization (Kidney, Uniform Negatives) | **0.845** | 0.506 | 0.626 |
| Tissue Colocalization (Lung, Uniform Negatives) | **0.837** | 0.642 | 0.519 |
| Tissue Colocalization (Lymph Node, Uniform Negatives) | **0.803** | 0.646 | 0.545 |

Supervision lifts interaction off its cosine floor, 0.586 to 0.741 on the large-n split, and stops short of HI-JEPA at 0.950, with non-overlapping intervals. On the tasks that require measured proximity it does not close the gap: knockout recovery is unchanged (0.108 against 0.119 for cosine), and on tissue the probe helps in one organ and hurts in two. Supervision cannot recover a signal the sequence does not carry.

### 11.2 Tissue Colocalization by Organ

This task ranks pairs of proteins by whether they sit close together within an organ’s tissue. It is proximity-grounded and is a weaker claim than physical binding, since two proteins can share a neighborhood without contact.

Abundant proteins colocalize with many partners, so the partner-count correction matters most at this scale: on kidney, a score built from partner counts alone reaches 0.707 against uniform negatives and 0.503 once they are matched. The model leads ESM-C in every organ under matched negatives.

## 12. Discussion and Scope

**What the paper establishes.** A representation trained on measured molecular proximity performs operations that sequence and structure models cannot, connects scales in one coordinate system, and recovers the established mediators of amyloid pathology from tissue in which no disease label was supplied. The ablation of Section 9 shows that the cross-scale operation comes from the proximity measurement and from nothing else in the training signal.

**Where the scope ends.** Three boundaries.

1. Disease results contrast diseased tissue against control within an organ, and cover brain and liver (§8.2). How a configuration remodels from one disease state to another is not addressed.
2. Co-assembly recovery within a tissue is a broader check than the organ-level colocalization of Table 9, which is scored on five organs. It holds across seven, and the eighth is too noisy to score.
3. The forward transition under treatment is a prospective prediction registered here, not a measured result. Validation of the trained encoder on human ExM tissue is future wetlab work.

**Table 9.** Tissue-Scale Colocalization. Shipped checkpoint. A pair counts as a positive when its colocalization is statistically significant in at least three held-out samples of that organ, which excludes pairs appearing in a single sample by chance. Negatives are matched to the positives on how many partners each protein colocalizes with across the corpus; values are the mean over eight draws. The last column scores a pair using nothing but those two counts.

| Organ | Pairs (n) | HI-JEPA | ESM-C 6B | Partner Count |
| --- | --- | --- | --- | --- |
| Colon | 29 | **0.859** | 0.254 | 0.621 |
| Cardiac Atrium | 66 | **0.847** | 0.492 | 0.493 |
| Lung | 657 | **0.811** | 0.609 | 0.504 |
| Kidney | 829 | **0.796** | 0.557 | 0.503 |
| Lymph Node | 329 | **0.777** | 0.634 | 0.502 |

**Against virtual-cell programs.** A virtual cell simulates the states of a single cell, where HI-JEPA models the measured physical organization of molecules from molecular assembly through tissue to disease. Virtual-cell proposals use generative encoder–decoder-and-transition designs [12], where HI-JEPA predicts in latent space, never reconstructs its input, and learns from measured positives alone (§5).

**What distinguishes the model is its input.** Measured molecular proximity at nanoscale is a signal that models trained on sequence, structure, annotation or expression do not have, and §9 shows that removing it is what costs the cross-scale operation. Multiplex spatial imaging extends it at coarser resolution for breadth and human translation. Expanding that corpus is the principal avenue for improvement, and measurement throughput is what limits it, not compute. Throughput is bounded by how many proteins one acquisition can resolve, since conventionally each target requires its own labeled reagent. Reading identity from geometry (§6.2) removes the per-target reagent from that chain, so the instrument and the model develop in tandem: the model improves as the corpus grows, and the corpus grows on the proteins the model can already identify.

**On priority.** To our knowledge no prior model of biology takes measured molecular proximity in intact tissue as the state it perturbs and plans over. We state it in that bounded form rather than as a claim to the first “world model of biology”, because the term is contested (§1.3) and the claim would rest on a definition rather than on a result.

## 13. Conclusion

We built a joint-embedding predictive model of biology whose state is measured molecular proximity. It observes what a sequence model cannot read, acts on a configuration that can be perturbed, and plans interventions scored along an axis fit from real diseased and healthy tissue. The objective learns from measured positives alone, leads a contrastive control trained on identical data across every capability, and reaches comparable interaction AUC at roughly 2,000 times fewer pair comparisons without its dimensional collapse.

Ablation shows that the operation connecting the scales comes from the proximity measurement and from nothing else in the training signal. That measurement is ours to extend, and identifying proteins from geometry rather than from reagents is what lets us extend it.

## Availability

This paper reports capabilities. The training procedure, the data pipeline and the measurement-acquisition specifics are proprietary and are not described. Section 4.2 gives the evaluation protocol behind every number reported. The proximity measurement is reproducible on public data: the 5xFAD study reanalyzes a public deposit, so a reader can obtain the same images and check how an acquisition becomes a measured relation. Benchmark splits and derived evaluation data can be released; code, weights and the proprietary corpus are not. Every number reported here is tied to a stored result and re-derived from it before each build.

## Broader Impacts

A pocket that opens only in a pathological configuration, or an interface that forms only between proteins a disease state has brought together, is invisible to a method reading sequence or a single structure. Targets of that kind are what this model is built to find, and measuring where molecules sit in diseased tissue is what puts them in reach. Small molecules, peptides, molecular glues and degraders can all be directed at them once they are.

The conditions this is aimed at are ones where conventional target discovery has repeatedly failed, neurodegeneration and chronic pain among them [26]. What the model produces is a target nomination and a predicted consequence, and each nomination is tested in our own wet lab.

